## Supplementary material for "Axial variation of deoxyhemoglobin density as a source of the low-frequency time lag structure in blood oxygenation level-dependent signals": Supp. Fig.

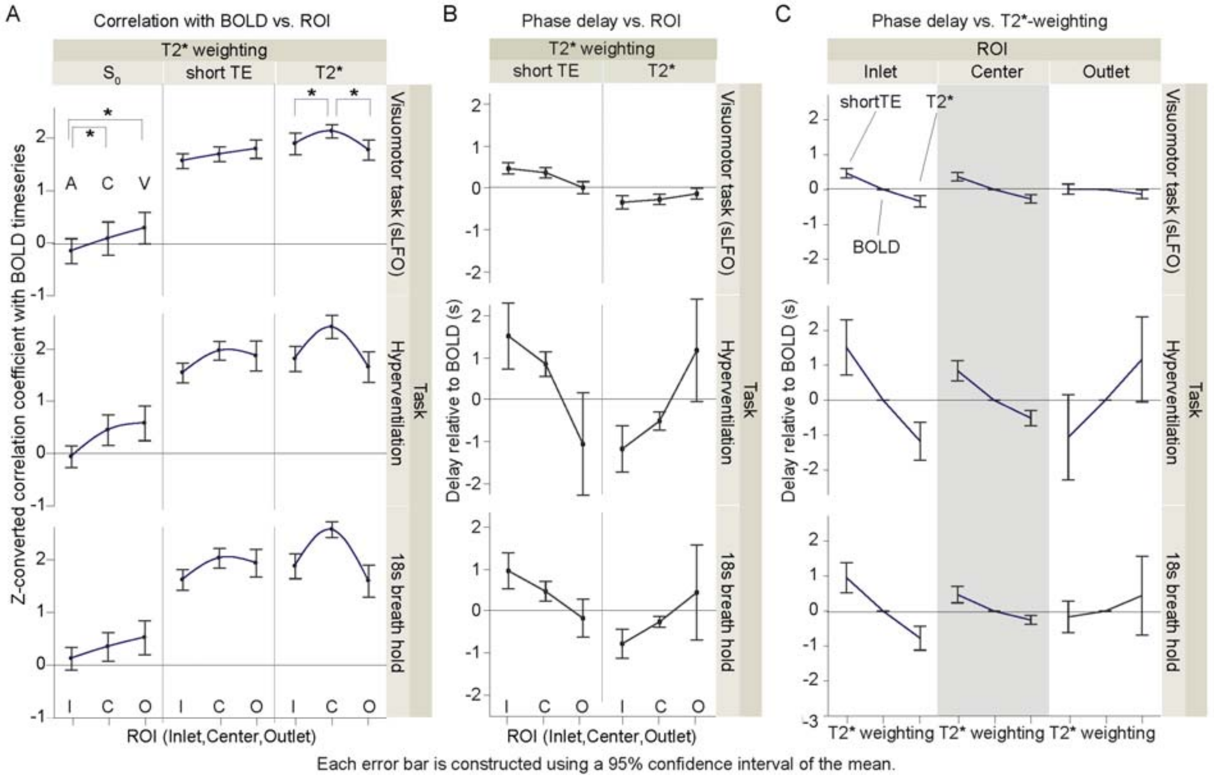


**Supplementary Figure 1. A,** Correlation analysis of the multi echo-derived signals extracted from the three vascular regions. Effects of vascular region was observed on both $S_0$ and T2* components, but in a different manner. **B,** Phase analysis of the low-frequency component below 0.1 Hz. Phase delay relative to BOLD time course was calculated for the two T2* weighted signals. Respiratory challenges enhanced the phase difference between $S_0$ and T2*. **C.** The same data as in B, but separately plotted for each region. Dissociation of the T2* and $S_0$ phase, as well as its interaction with the vascular region is evident. All these effects were small with spontaneous low-frequency fluctuation but enhanced in artificial oscillation by respiratory challenges.

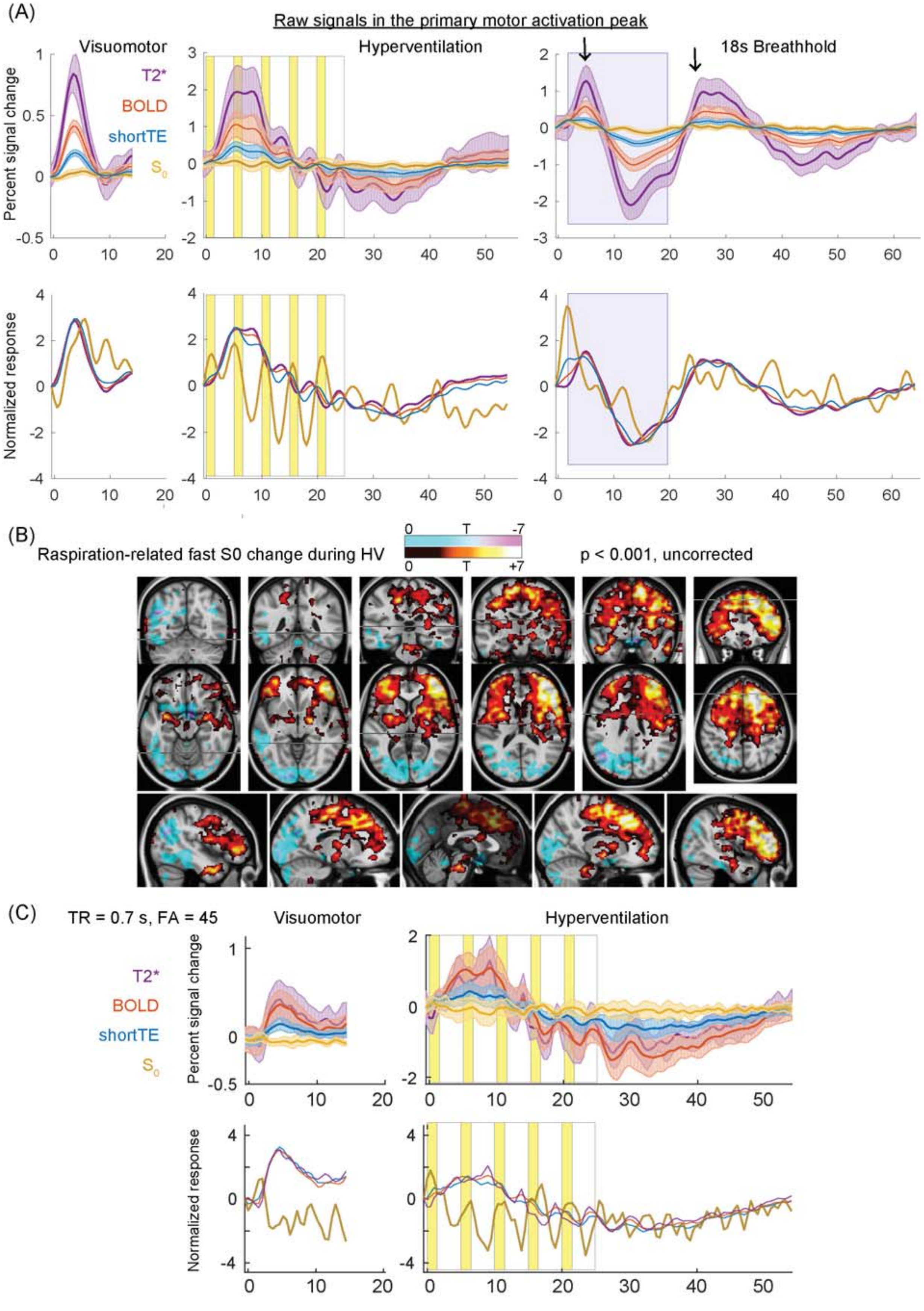
(A)
Raw signals in the primary motor activation peak
Visuomotor
Hyperventilation
18s Breathhold
T2*
BOLD
shortTE
$S_0$
Percent signal change
Normalized response
(B)
Raspiration-related fast S0 change during HV
p < 0.001, uncorrected
0 T -7
0 T +7
(C)
TR = 0.7 s, FA = 45
Visuomotor
Hyperventilation
T2*
BOLD
shortTE
$S_0$
Percent signal change
Normalized response

**Supplementary Figure 2. A**, Raw signal responses in the motor/premotor activation cluster for each task from Experiment 2. Time courses were resampled to a sampling interval of 0.5 s. Small but distinct responses were found in the non-BOLD or $S_0$ component but with a dominating fast component compared to the regional response, apparently independent of both the neurovascular coupling and hemoglobin fluctuations that accompany T2* changes. This response was roughly out of phase with the beat-to-beat mean arterial blood pressure shown in Fig. 3, although the blood pressure response was absent during the first two cycles. Arrows indicate the fluctuation corresponding to the neurovascular coupling modelled in the SPM analysis. **B.** Group SPM to assess the distribution of the respiratory phase-related fast fluctuation of $S_0$ during hyperventilation. A separate SPM analysis was performed by modeling the fast $S_0$ fluctuation only. An anterior-posterior segmentation is evident. In addition, there are small symmetrical structures along the major veins. **C.** Additional experiment from a subset of participants (N = 7) using a different set of repetition time/flip angle to manipulate the T1-inflow effect on the $S_0$ component. The overall $S_0$ response is lower than those shown in panel **A**, but the fast response to hyperventilation is preserved.
